# A MICROTUBULE ASSOCIATED PROTEIN is required for division plane orientation during 3D-differential growth within a tissue

**DOI:** 10.64898/2026.01.07.697928

**Authors:** Zsófia Winter, Dorothee Stöckle, Takema Sasaki, Sophie Marc Martin, Yoshihisa Oda, Joop EM Vermeer

## Abstract

Communication between tissues and cell division orientation are essential for organ development. During lateral root formation in *Arabidopsis thaliana,* three-dimensional differential growth is established within a tissue of interconnected and pressurized cells. To emerge, the new organ needs to overcome the mechanical constraints provided by the overlying tissues. How the division plane orientation within the growing organ encountering mechanical constrains is coordinated remains unclear.

Here, we show that MICROTUBULE ASSOCIATED PROTEIN 70-2 contributes to this integration of mechanical feedback during lateral root development by being associated with division plane orientation during primordium formation. MAP70-2 expression in the lateral root primordium is dynamic, localizing to cell corners and the cortical division zone prior to cytokinesis. Loss of *MAP70-2* leads to misoriented cell divisions in the early stage of lateral root development and defective morphogenesis. Here we propose that MAP70-2 integrates biochemical and mechanical cues to establish correct division plane orientation during three-dimensional differential growth to facilitate lateral root organogenesis.

## Introduction

In plant development, the orientation of the division plane is a critical determinant of cell fate, tissue patterning, and organ morphogenesis. Division plane orientation is established by cortical markers put in place by the preprophase band (PPB), which is required for the expanding phragmoplast to the correct cortical site. This spatial coordination ensures accurate cell plate insertion, although the molecular mechanisms linking division site specification to phragmoplast guidance remain only partially understood [1–3]. The proper establishment of the cortical division zone (CDZ) is essential in plant cells, as it precisely determines where cells will divide to establish their body plan. The CDZ, initially marked by the PBB forming at the G2 phase of mitosis, serves as a guide for the phragmoplast, which deposits the new cell plate during cytokinesis [4,5]. Accurate positioning of the CDZ ensures that daughter cells are correctly oriented, maintaining organized tissue architecture and contributing to proper organ development [6]. Several proteins are essential for establishing and maintaining the CDZ. The **T**ONNEAU – **T**ON-RECRUITING-MOTIF – **P**ROTEIN PHOSPHATASE 2A (TTP) complex is required for PPB formation, acting through microtubule organization at the cortex [7–9]. *MICROTUBULE ORGANIZATION 1* (*MOR1*), a microtubule polymerase, and CLIP-ASSOCIATED PROTEIN (CLASP), a microtubule plus-end stabilizer, also play vital roles in PPB formation and microtubule dynamics [10,11]. Once the PPB disassembles, CDZ positional memory is maintained by proteins like TANGLED1 (TAN1), RANGAP1, PHRAGMOPLAST ORIENTING KINESIN 1 and-2 (POK1/POK2) that guide the phragmoplast to the correct cortical site [6,12–15]. IQ DOMAIN (IQD) proteins and PLECKSTRIN HOMOLOGY RHO GTPASE ACTIVATING PROTEINS (PHGAPs) further contribute to CDZ stability by linking cortical signals to the cytoskeleton [16,17]. This system is tightly regulated by phosphorylation events mediated by AURORA kinases (AUR1/2), which interact with microtubule-associated proteins like MICROTUBULE ASSOCIATED PROTEIN 65-3 (MAP65-3) to control phragmoplast dynamics [18]. Improper CDZ formation can result in misplaced division planes, disrupted asymmetric divisions, and mechanical discontinuities due to misaligned cell walls [6,19]. Thus, the CDZ serves as a physical location marking cell division orientation, essential for proper organogenesis.

The organogenesis of lateral roots (LRs) allows the exploration of the soil to proliferate into nutrient-rich regions. In *Arabidopsis thaliana* (Arabidopsis), LRs initiate from the xylem pole pericycle (XPP), located deep within the differentiation zone of the primary root between the xylem and endodermis [20,21]. Arabidopsis LRs originate from specialized XPP cells, known as LR founder cells (LRFCs). One of the defining features of LR initiation in Arabidopsis is the asymmetric swelling of LRFCs, accompanied by the directed movement of their nuclei towards the shared cell wall. This is followed by a formative cell division, a process that relies heavily on the precise reorientation of the division plane, resulting in a stage I LR primordium. Proper control of cell morphology and division plane alignment during LR initiation depends on the coordinated integration of both cell-and non-cell-autonomous signals [22,23]. In Arabidopsis, LR development can be divided into eight stages (Stage I-VIII). During this process, the LR needs to grow through the endodermis, cortex, and epidermis cell layers to emerge at the surface [20,24]. LR emergence is a biphasic process: in the first morphogenetic phase (Stage I-IV), a four-layered LRP is formed in the XPP that is confined by the endodermis. The second phase consists of three stages in which a dome-shaped meristem structure is created, traversing of the endodermis that is followed by rapid organ growth through the remaining two overlying cell layers [20,24,25].

LR development is tightly regulated by auxin signaling, which coordinates both founder cell specification and the remodeling of overlying tissues to allow organ outgrowth. During this process, endodermal cells overlying the primordium, adapt their shape through a regulated loss of cell volume triggered by Aux/IAA mediated signaling in these cells. Blocking this nuclear auxin signaling pathway in the differentiated endodermis blocks LR formation [26]. However, it is still unclear how LRFCs and the neighboring endodermis communicate in order to channel organ growth via spatial accommodating responses. In addition, besides the role of AURORA kinases and MAP65 proteins, little is known how the correct cell division planes are established in the primordium during its organogenesis.

Plant cortical microtubules (CMTs) are nucleated at the cell cortex and remain linked to the plasma membrane throughout the nonmitotic phases of the cell cycle [12]. The transversally co-aligned CMTs enable anisotropic cell growth in hypocotyls, stems and roots [13]. Microtubule-associated proteins (MAPs) have been implicated in various organogenic processes, including the directional growth of petals, the uneven expansion of pavement cells in the Arabidopsis leaf epidermis, and the regulation of fruit initiation and size in tomato [27–29]. A plant-specific MAP family with an approximate size of 70 kDa, the MAP70s, comprises 5 members in Arabidopsis and is distinct in that it shows no sequence homology with other known MAPs [30]. Among its members, *MAP70-1* and *MAP70-5* are required for cell wall remodeling and directional growth, and both proteins have been observed to co-sediment with microtubules *in vitro* [31]. In 7-day-old Arabidopsis seedlings, *MAP70-1* and *MAP70-5* were shown to play a role in guiding the formation of cell wall ingrowths at the edges of xylem pits [32]. In addition, *MAP70-5* has been reported to reduce microtubule stiffness and is necessary for CMT reorganization in the overlying endodermis during LR development [32].

Here, we show that another member of the MAP70 family, *MAP70-2* is expressed the root apical meristem and in primordia throughout LR development. MAP70-2 displays a dynamic localization that appears to predict cell division plane orientation and its expression domain in the primordium is correlated with meristem establishment. Loss-of-function mutants of *MAP70-2* display shorter primary roots, and abnormal LR development. The latter appears to be due to problems in the orientation of the cell division plane, which is specific for LR development. MAP70-2 function appears to be distinct from that of MAP70-5 and knocking out *MAP70-2* in the *map70-5-c1* mutant [33] restores normal LR formation. This suggests that both proteins have specialized roles in maintaining an organ-specific mechanostasis during organ development.

## Results

### MAP70-2 localizes to cortical domains and future division planes in root meristems

To characterize the expression of *MAP70-2* in seedlings we used plants transformed with a *MAP70-2pro::NLS-EGFP:GUS* reporter (Figure 1A). This revealed that *MAP70-2* was expressed in root meristems, but also around the vascular system of the cotyledons (Figure 1A). To investigate the subcellular localization and dynamics of MAP70-2 during LR development, we imaged Arabidopsis roots co-expressing *MAP70-2pro::CITRINE:MAP70-2* and a membrane marker, *LBD16pro::3xmCherry:SYP122*, from initiation to near emergence LR (Stages I–VII) (Figure 1E). In stage I-II primordia, CITRINE:MAP70-2 was mostly localized to the cytoplasm but also co-localized with the plasma membrane marker at the cell edges, where it progressively accumulated at sites coinciding with future cell divisions. As the cell cycle advanced, CITRINE:MAP70-2 also appeared to localize in the corners of cells prior to cell division (Figure 1E; Stage II and Stage IV). From stage IV and beyond, CITRINE:MAP70-2 appeared to be localize along the traverse cell periphery of newly divided cells in the center of the primordium, suggesting a potential role in reinforcing the orientation and integrity of division planes during LR morphogenesis (Figure 1E; Stage V and Stage VI). In addition, the expression domain of *MAP70-2pro::CITRINE:MAP70-2* changed as LR development progressed. From stages I to III, CITRINE:MAP70-2 fluorescence could be detected throughout the primordium, whereas from stage IV, CITRINE:MAP70-2 appeared to accumulate in the central part of the primordium. Moreover, the signal of CITRINE:MAP70-2 appeared to be the strongest in the central cells of the primordia, most likely reflecting the establishment of a new meristematic region in the new organ (Figure 1E). To better visualize the dynamics of CITRINE:MAP70-2, we used 4-dimensional (4D) live cell 2-photon microscopy to follow its localization during LR development. This revealed the dynamics of CITRINE:MAP70-2 throughout the cell divisions and the shift of its expression domain towards the central part of the primordium. The latter also was evident when observing the emergence of a LR, where CITRINE:MAP70-2 showed a more pronounced signal in the meristem of the LR (Supplemental Movies S1-2).

**Figure 1.**
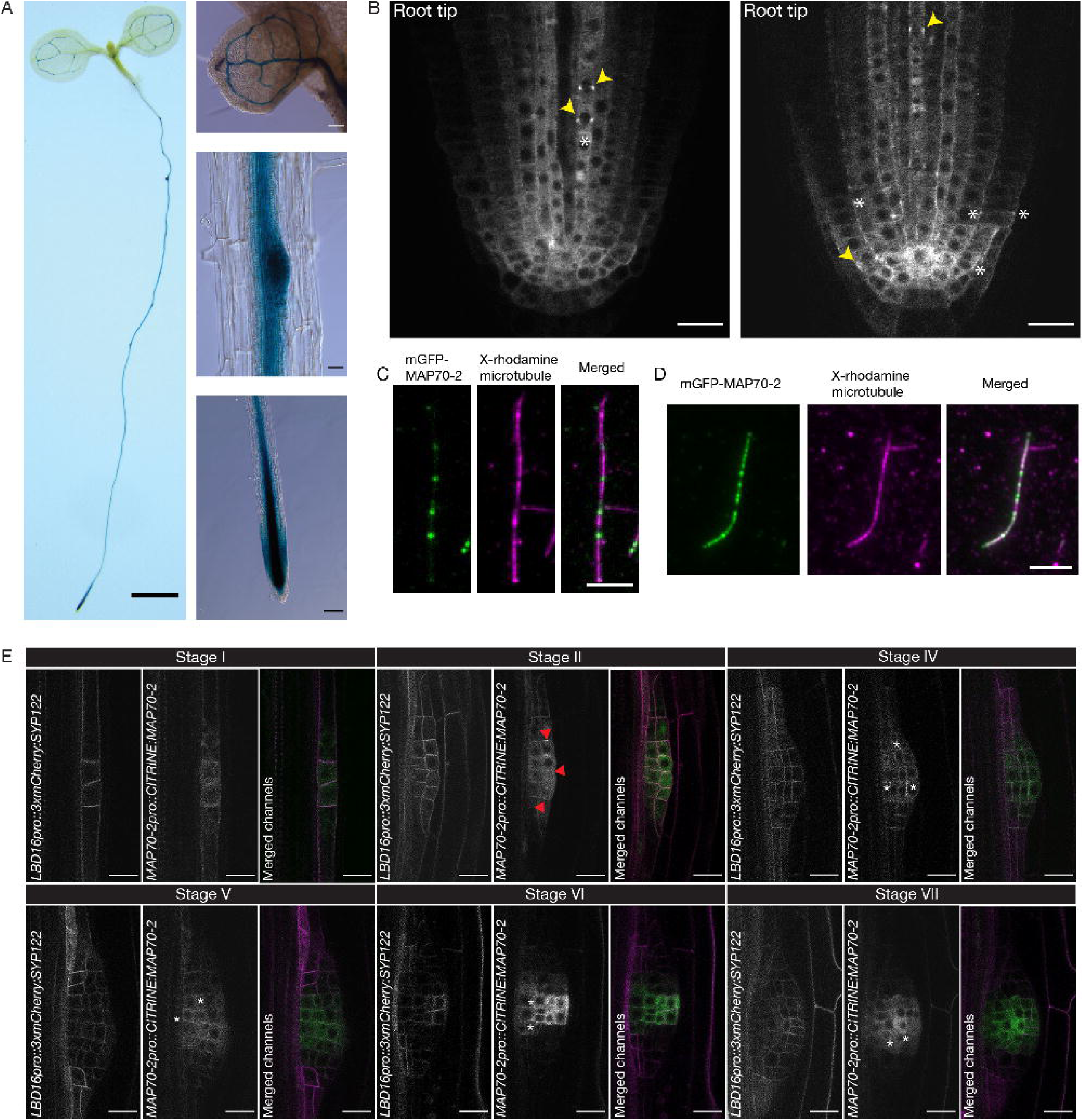
MAP70-2 displays a dynamic localization pattern in root meristems and binds microtubules *in vitro*. (A) GUS staining patterns in 5-day-old transgenic Arabidopsis (Col-0) seedling carrying the *MAP70-2pro::NLS-GFP:GUS* transgene. *MAP70-2* shows expression in plant vasculature, lateral root primordia and primary root apical meristem. (B) Confocal images of two different z-planes of the same root tip expressing CITRINE:MAP70-2 showing localization to the cell division zone (yellow arrow heads), as well as lateral/corner accumulation (white asterisks). (C and D) Purified mGFP:MAP70-2 localizes to *in vitro* polymerized, rhodamine-labeled microtubules. (C) 5nM mGFP-MAP70-2 and (D) 15 nM mGFP:MAP70-2. (E) CIRINE:MAP70-2 localization dynamics during early stages of LR development. Stage I primordia already express *MAP70-2pro::CITRINE:MAP70-2* (green). From stage II to Stage VII, CITRINE:MAP70-2 accumulates in the apical and central cells of the developing LR primordium. Red arrowheads show CITRINE:MAP70-2 accumulation at the cell division sites and white asterisks indicate corner localization of CITRINE:MAP70-2. The plasma membrane is visualized by the LBD16pro::3xmCherry:SYP122 marker (magenta). For each developmental stage at least 10 roots in 3 independent experiments were analyzed. Scale in (A) whole seedling = 1mm; hypocotyl, primary root tip scale bar = 100 µm. Scale in (B) = 20 µm, in (C and D) = 5 µm, (E) in = 10 µm.

Consistent with these results, imaging of primary root tips expressing *MAP70-2pro::CITRINE:MAP70-2* revealed similar localization patterns (Figure 1B). In contrast to CITRINE:MAP70-2 localization in primordia, in the root apical meristem, CITRINE:MAP70-2 was also localizing to filamentous structures resembling cortical microtubules (CMTs) (Figure 1B). In general, CITRINE:MAP70-2 was broadly distributed along the cortical sides of cells in the meristematic zone, but showed distinct enrichment at transverse walls of recently divided or dividing cells, suggesting an association with sites of cell division. Notably, some cells displayed MAP70-2 polarization along presumptive division planes prior to visible cytokinesis, suggestive of a role in division site establishment (Figure 1B). Depolymerization of microtubules by oryzalin revealed that the localization of CITRINE:MAP70-2 required intact microtubules as the treatment resulted in a diffuse cytosolic localization both in the root apical meristem and the LR primordium (Supplemental Figure S3A and B). To confirm whether MAP70-2 could directly bind microtubules, in vitro purified mGFP:MAP70-2 protein was incubated with rhodamine labelled, in vitro polymerised microtubules, which revealed that mGFP:MAP70-2 could directly bind microtubules (Figure 1C and D). Together, these data suggest that MAP70-2 can bind microtubules directly and possibly functions in organizing or stabilizing CMT arrays that contribute to division plane orientation in both primary and LR development. Interestingly, although we did not observe labelling of microtubules by CITRINE:MAP70-2 in LR primodia, its localization to the cell periphery was sensitive to oryzalin treatment (Supplemental Figure 3A and B), suggesting a cell type dependent localization of MAP70-2.

### MAP70-2 is required for lateral root development

To assess the role of MAP70-2 in primary and LR development, we generated loss-of-function mutants carrying large deletions in the central domain that is predicted to contain the microtubule binding region (Supplemental Figure 1A) [32,34]. First, we compared root growth between wild-type Col-0 and two independent *map70-2* mutant alleles (*map70-2–c1* and *map70-2–c2*). This revealed that both *map70-2* alleles had a shorter root compared to wild type (Figure 2A and B). Quantification of the root meristem size revealed that the *map70-2* mutants had a reduced meristem (Figure 2C). Next, we quantified the number of LRs per centimeter of lateral root development zone (LRDZ) in the *map70-2* mutants. Both alleles exhibited a significantly higher number of LRs compared to Col-0 (p < 0.001), suggesting that normal progression of LR development requires a functional *MAP70-2* (Figure 2D). Analysis of the distribution of the different stages of LR development revealed that the observed increase in LRs in *map70-2* mutants appeared not due to enhanced initiation, as the number of stage I primordia was not affected, but rather due to a significant increase in stage II primordia (Figure 2E). Introgression of the *MAP70-2pro::CITRINE:MAP70-2* reporter rescued both the root length as well as the LR phenotype, indicating that the CITRINE:MAP70-2 fusion is functional (Supplemental Figure S2). Confocal imaging of LRs in *map70-2–c1* roots expressing the plasma membrane marker ENHANCED YELLOW FLUORESCENT PROTEIN:NOVEL PLANT SNARE 12 (EYFP:NPSN12; W131Y [35]) revealed distorted cellular organization and irregular division patterns during the transition from stage II to III during LR development (Figure 2F and G), indicating that MAP70-2 is required for proper morphogenetic transitions during LR development. It appears that MAP70-2 is required to facilitate the progression of primordia beyond Stage II, possibly by integrating mechanical constraints imposed by the surrounding tissue [33,36–38].

**Figure 2.**
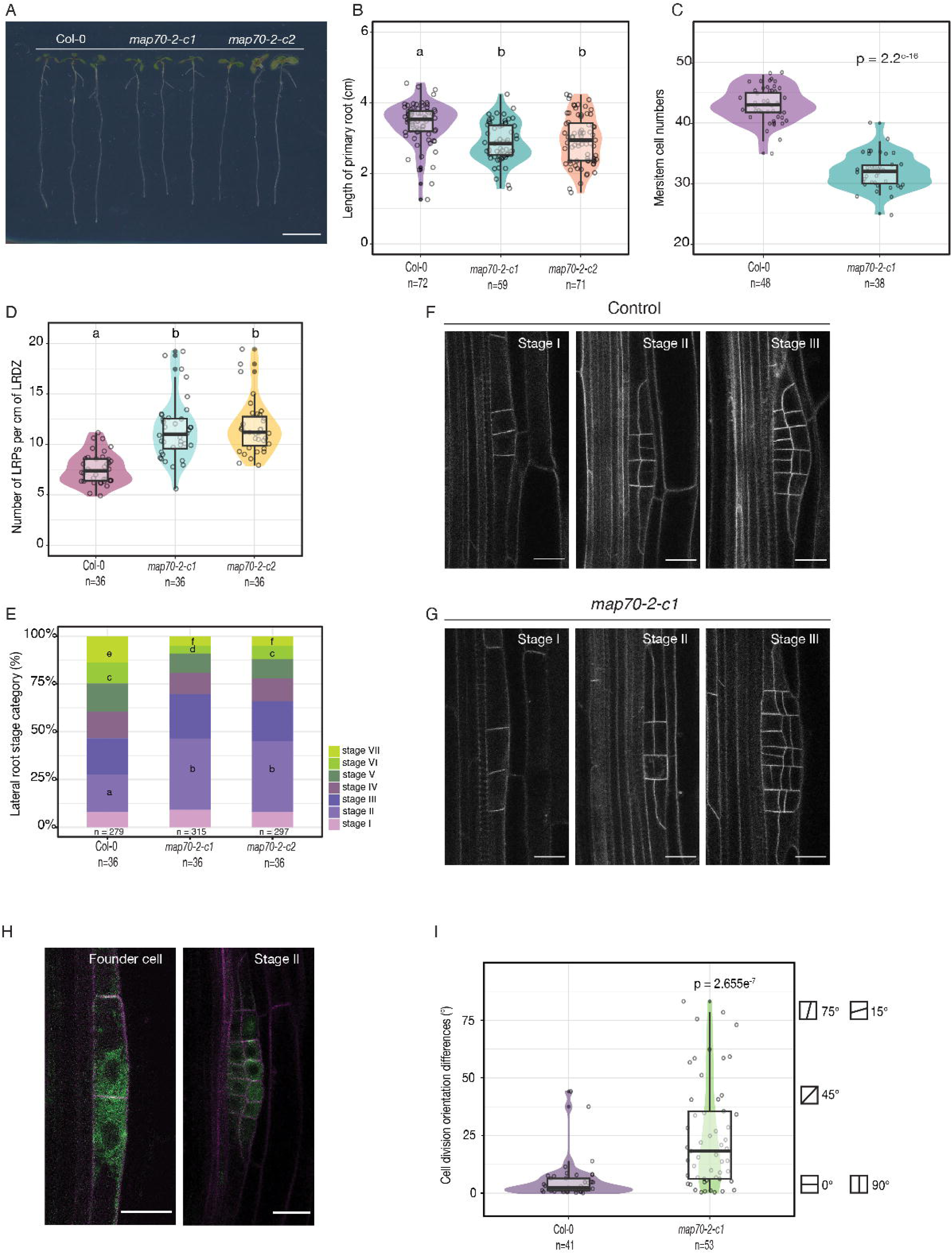
MAP70-2 is required for lateral root development. (A) Image of 7-day-old wild type (Col-0), *map70-2-c1* and *map70-2-c2* seedlings showing that MAP70-2 is required for root growth. (B) Quantification of the total root length. Primary roots of both *map70-2* alleles are significantly shorter than Col-0. Two separate student‘s t-tests on Col-0 and *map70-2-c1* and *map70-2-c2* were performed. (*map70-2-c1* *p= 0,00019, *map70-2-c2* *p= 0.0002). (C) Comparing meristem cell numbers between Col-0) and *map70-2-c1*. The difference between Col-0 and *map70-2-c1* was assessed using an unpaired two-sample t-test assuming equal variances. The meristem cell counts are significantly reduced in *map70-2-c1* compared to Col-0, as indicated by a highly significant p-value (p = 2.2e-16). Sample sizes are shown in the graphs as n. (D) Comparison of the LR density in the LRDZ of Col-0 compared to *map70-2-c1* and-*c2*. Wilcoxon rank-sum test with continuity correction was performed for each line. (*map70-2-c1* p= 2.66e-8, *map70-2-c2* p= 1.961e-9). (E) Staging of LR development revealed a significant higher number of stage II and stage III LRP in both *map70-2* alleles compared to Col-0. LRP stage distribution on each genotype was assessed by comparing stage frequencies between Col-0 and each mutant allele using Pearsońs Chi-squared test with Yateś continuity correction (*map70-2-c1* stage II= 6.4e-5, stage VI= 0.0073, stage VII= 0,00095, *map70-2-c2* stage II= 3.12e-5, stage VII= 0.003). (F and G) LRP morphology in WT and *map70-2-c1* through stages I-III visualized by the plasma membrane marker *UBQ10pro::EYFP:NPSN12.* The overall percentage of deformed LRPs was significantly higher in the *map70-2-c1* mutant (64.4% of n=45) compared to wild-type Col-0 (31.25% of n=48). (H) Images of LR founder cells and stage Il LRPs expressing *LBD16pro::3xmCherry:SYP122* (plasma membrane, magenta) to visualize the LRP and *MAP70-2pro::CITRINE: MAP70-2* (green). White arrowheads show accumulation of CITRINE:MAP70-2 in the division site, both anticlinal (founder cell) and periclinal (stage II). (I) Quantification of forming cell plate angles in dividing LRP cells (stage I-III LRP) relative to the growth axis in the Col-0 and in *map70-2-c1*. Significance was determined based on two-sample Student‘s t-tests with a p-value = 2.655e-7. The violin plots show the distribution of division angles, where the shape of the violins outlines the density. The central lines show the medians. Scale bar in (A) 1 cm, (F – H) 20 μm.

### MAP70-2 is required for cell division plane orientation in lateral root primordia

In Arabidopsis lateral root founder cells, CMT reorientation is required to enable the asymmetric, radial expansion that precedes formative divisions [3,33,38]. Several MAPs have already been identified as key regulators of cytoskeletal organization and division plane establishment, underscoring their role in coordinating morphogenesis [13,16,31,32]. Based on the observed abnormalities in the cellular organization in stage II LR primordia in *map70-2-c1*, we quantified cell division plane orientation by comparing the angles of division planes in developing LRPs. This revealed significant differences between Col-0 and the *map70-2-c1* mutant. As shown in Figure 2H, MAP70-2 is recruited to the cell cortex, consistent with its potential involvement in both anticlinal and periclinal divisions. To visualize and quantify microtubule organization in developing LRPs, we generated an *LBD16pro::TurboID(TbID)-SYFP1:MBD* reporter line in both Col-0 and *map70-2-c1* backgrounds, enabling cell type-specific labeling of CMTs. This allowed direct comparison of the cell plate positioning angles between the two genotypes. This (Figure 2I) revealed that *map70-2-c1* LRPs exhibited a significantly more variable range of division angles compared with Col-0 (p = 2.655 × 10⁻⁷; between Col-0 and *map70-2-c1*). These results indicate that MAP70-2 function appears critical for maintaining proper cell division orientation during early LR development, potentially influencing primordium patterning and organogenesis. Interestingly, when comparing the organization of the primary root meristem between Col-0 and *map70-2-c1*, we did not observe any abnormalities in the organization of the cell files, suggesting that the role of MAP70-2 in regulating cell division plane orientation is specific for the LR development (Supplemental Figure S3C and D).

### MAP70-5 can partially replace MAP70-2

The Arabidopsis MAP70 family is divided into two clades: the MAP70-1 clade, which includes MAP70-2, and the divergent, eponymous MAP70-5 clade [34]. However, they show related localization patterns, as both MAP70-1 and MAP70-5 have been reported to decorate CMTs and regulate their organization during secondary cell wall deposition [32,34]. This suggests that despite their phylogenetic separation, these isoforms appear functionally connected [30,34]. Although they have different expression domains during LR development, both MAP70-2 and MAP70-5 can associate with CMTs, suggesting potential shared biochemical functions within the MAP70 family (Supplemental Figure 3A and B; and [33]). Both MAP70-2 and MAP70-5 are expressed in tissues, cells in the developing LRP or the overlying endodermis undergoing spatial accommodation, where CMT organization underlies key developmental processes (Figure 1E, [33,38]). This led us to investigate whether MAP70-5 could functionally replace MAP70-2, since both are required for LRP development, albeit in different tissues. Therefore, we expressed *TbID-SYFP1:MAP70-5* under the control of the *MAP70-2* promoter (*MAP70-2pro::TbID-SYFP1:MAP70-5*) and examined its ability to rescue the root phenotypes of the *map70-2-c1* mutant. The introgression of *MAP70-2pro::CITRINE:MAP70-2* reporter in *map70-2-c1* fully complemented the primary root length and LRP phenotypes in 7-day-old seedlings (Supplemental Figure 2), suggesting that N-terminal protein fusions are functional. In contrast, introgression of the *MAP70-2pro::TbID-SYFP1:MAP70-5* fusion revealed only a partial complementation. LRP density was restored to wild-type levels, but both *MAP70-2pro::TbID-SYFP1:MAP70-5* lines retained an elevated number of stage II primordia compared to Col-0, but lower compared to *map70-2-c1* (Figure 3C). The observation that theTbID-SYFP1:MAP70-5 fusion under the control of the MAP70-2 promoter appeared mostly cytosolic could be a possible explanation for the partial complementation (Figure 3A). Thus, it appears that the progression from stage I to stage III primordia is *MAP70-2* dependent. When analyzing the effect of primary root growth restoration, the *MAP70-2pro::TbID-SYFP1:MAP70-5* lines had slightly, but significant longer roots than Col-0 (Figure 3B). This suggests that MAP70-5 can replace MAP70-2 during primary root development.

**Figure 3.**
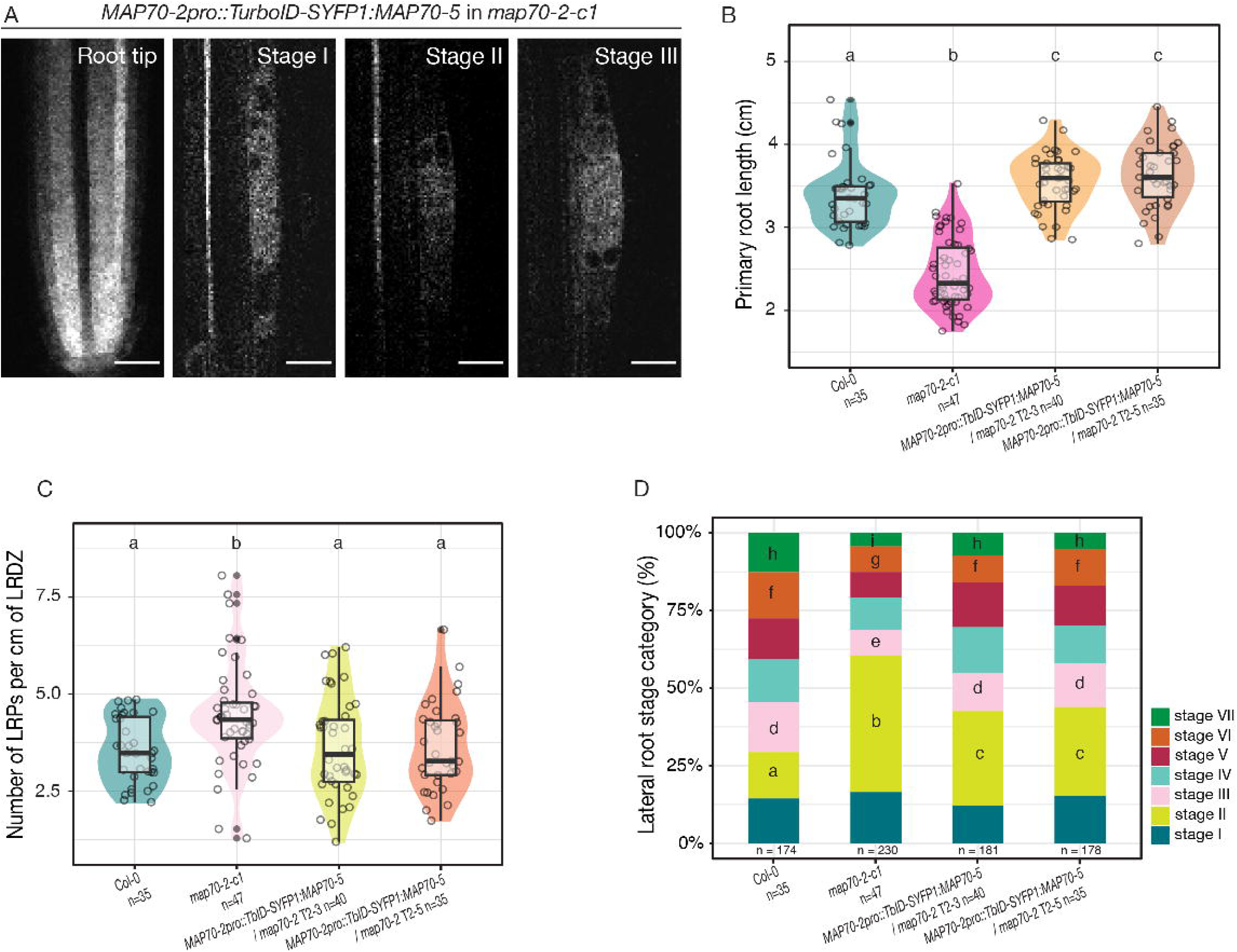
*MAP70-5* can partially functionally replace *MAP70-2*. (A) Confocal images showing the localization of CITRINE:MAP70-5 under the control of the *MAP70-2* promoter (*MAP70-2pro:TurboID-sYFP:MAP70-5*) in *map70-2-c1* roots. Representative images are shown for the root tip and for Stage I, Stage II, and Stage III of LR primordium development. Scale bars = 20 µm. (B) Comparison of primary root length between Col-0 and two independent *MAP70-2pro:TurboID-sYFP:MAP70-5* lines. Different letters above the plots denote statistically significant differences among groups (p < 0.05), as determined by appropriate Kruskal-Wallis rank sum test comparisons followed by Wilcoxon rank-sum test. (C) Comparison of LRP density in the LR developmental zone. Pairwise Wilcoxon rank-sum tests with Benjamini-Hochberg correction, P < 0.05 sum test was used to determine significant differences (p<0.05). (D) Comparison of the distribution of LR developmental stages. Total numbers of scored primordia (n) are indicated below each bar, and the number of seedlings analyzed per genotype is shown beneath genotype labels. Different letters within each stage denote statistically significant differences among genotypes (Pearson’s Chi-squared test with Yates’ continuity correction was used to assess if LRP distribution is different in promoter swapped transgenic lines, p < 0.05)

Remarkably, the *map70-2-c1/map70-5-c1* double mutant displayed overall root LR developmental stages comparable to wild type. However, *map70-2-c1/map70-5-c1* still exhibited a significantly shorter primary root with a reduced meristem and increased LR density (Figure 4E and C). Using the plasma membrane marker W131Y to compare LR morphology, no differences were observed between *map70-2-c1/map70-5-c1* and Col-0 (Figure 4F and G). These results, together with the previously described phenotypes of map70-5-c1 [33]suggest that *MAP70-2* and *MAP70-5* appear to regulate primary root development via different, non-additive mechanisms. In addition, it appears that the observed LR phenotypes observed in the *map70-2-c1* and *map70-5-c1* single mutants are dependent on the presence of a functional *MAP70-5* or *MAP70-2* gene in the neighboring cells.

**Figure 4.**
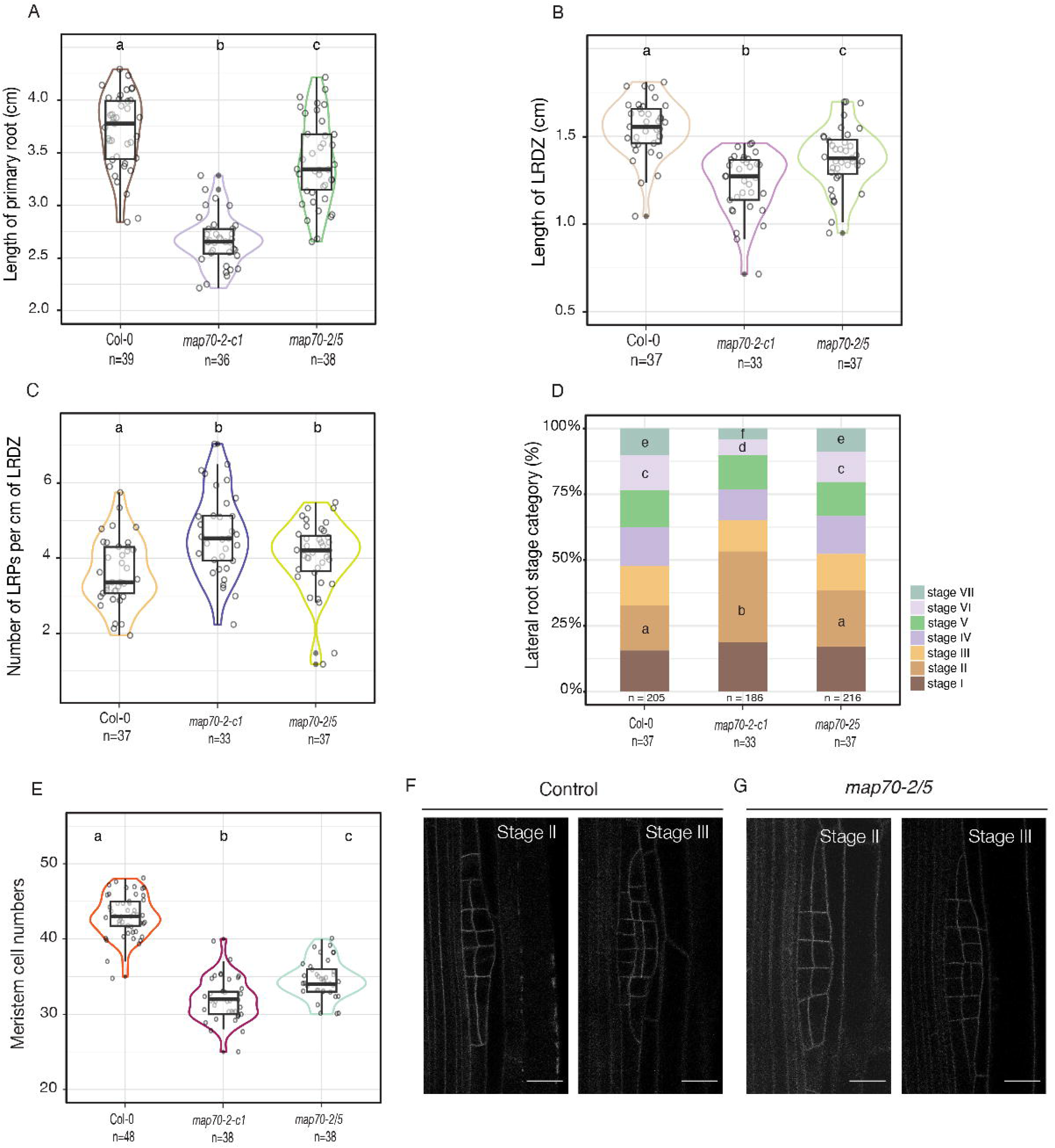
*MAP70-2* and *MAP70-5* both are required for normal primary root growth and lateral root development. (A) Comparison of primary root length between Col-0, *map70-2-c1* and *map70-2/5*. Statistical significance was assessed by one-way ANOVA followed by Tukey’s post hoc test (Col-0 vs. *map70-2/5*, p = 0.001349; *map70-2-c1* vs. *map70-2/5*, p < 0.001). (B) Comparison of LRDZ between Col-0, *map70-2-c1* and *map70-2/5*. Statistical significance among lines for quantification was assessed using the Kruskal–Wallis test followed by pairwise Wilcoxon rank-sum tests (Col-0 vs. *map70-2/5*, p = 1.9e-05; *map70-2-c1* vs. *map70-2/5*, p = 9.7e-04). (C) Comparison of LR density between Col-0, *map70-2-c1* and *map70-2/5*. Differences were evaluated using the Kruskal–Wallis test with post hoc Wilcoxon rank-sum comparisons (Col-0 vs. *map70-2/5*, p = 3.5e-04, *map70-2-c1* vs. *map70-2/5*, p = 0.0619). (D) Comparison of LR developmental stages between Col-0, *map70-2-c1* and *map70-2/5*. Numbers below the bar graph indicate total primordia and n below the genotype the amount of seedlings analyzed. Different letters denote significant differences (Pearson’s Chi-squared test with Yates’ continuity correction, p< 0.05). (E) Comparison of primary root meristem length between Col-0, *map70-2-c1* and *map70-2/5*. Statistical significance was assessed by one-way ANOVA followed by Tukey’s post hoc test (Col-0 vs. *map70-2/5*, p < 0.001; *map70-2-c1* vs. *map70-2/5*, p=7,12e-05). (F and G) Comparison of LRP morphology between Col-0 and *map70-2/5*. Stage II and III LRP were visualized using the plasma membrane marker *UBQ10pro::EYFP:NPSN12* (W131Y). The frequency of morphologically deformed primordia was comparable between the *map70-2/5* double mutant (52.17%, *n* = 45) and wild-type Col-0 (53.3%, *n* = 30).

### Overexpression of MAP70-2 causes twisted roots

To further explore the role of *MAP70-2* in root development, we generated *MAP70-2* overexpression lines using the *RPS5A* promoter (*RPS5Apro::TbID-SYFP1:MAP70-2*), which drives strong overexpression in dividing and expanding cells [39]. This approach allowed us to assess whether elevated *MAP70-2* levels could affect root development and CMT organization in actively growing tissues. The *RPS5Apro::TbID-SYFP1:MAP70-2* seedlings exhibited left-handed primary root growth and right-handed epidermal cell-file rotation (Figure 5A). Primary root length was comparable to wild type, meaning significantly longer than *map70-2-c1* (Figure 5B). By contrast, both overexpression lines had a shorter LRDZ, like *map70-2-c1* (Figure 5D). LRP density was indistinguishable from wild type in *RPS5Apro::TbID-SYFP1:MAP70-2 (2)*, whereas the other line showed an intermediate phenotype that was not significantly different from either wild type or *map70-2-c1* (Figure 5C, comparison to *map70-2-c1*: *p* = 0.005533). Notably, both lines accumulated elevated numbers of stage II LRPs relative to wild type, comparable to *map70-2-c1* (Figure 5E). These results indicate that the effects of MAP70-2 appear dose-dependent, as the overexpression lines show similar LR phenotypes.

**Figure 5.**
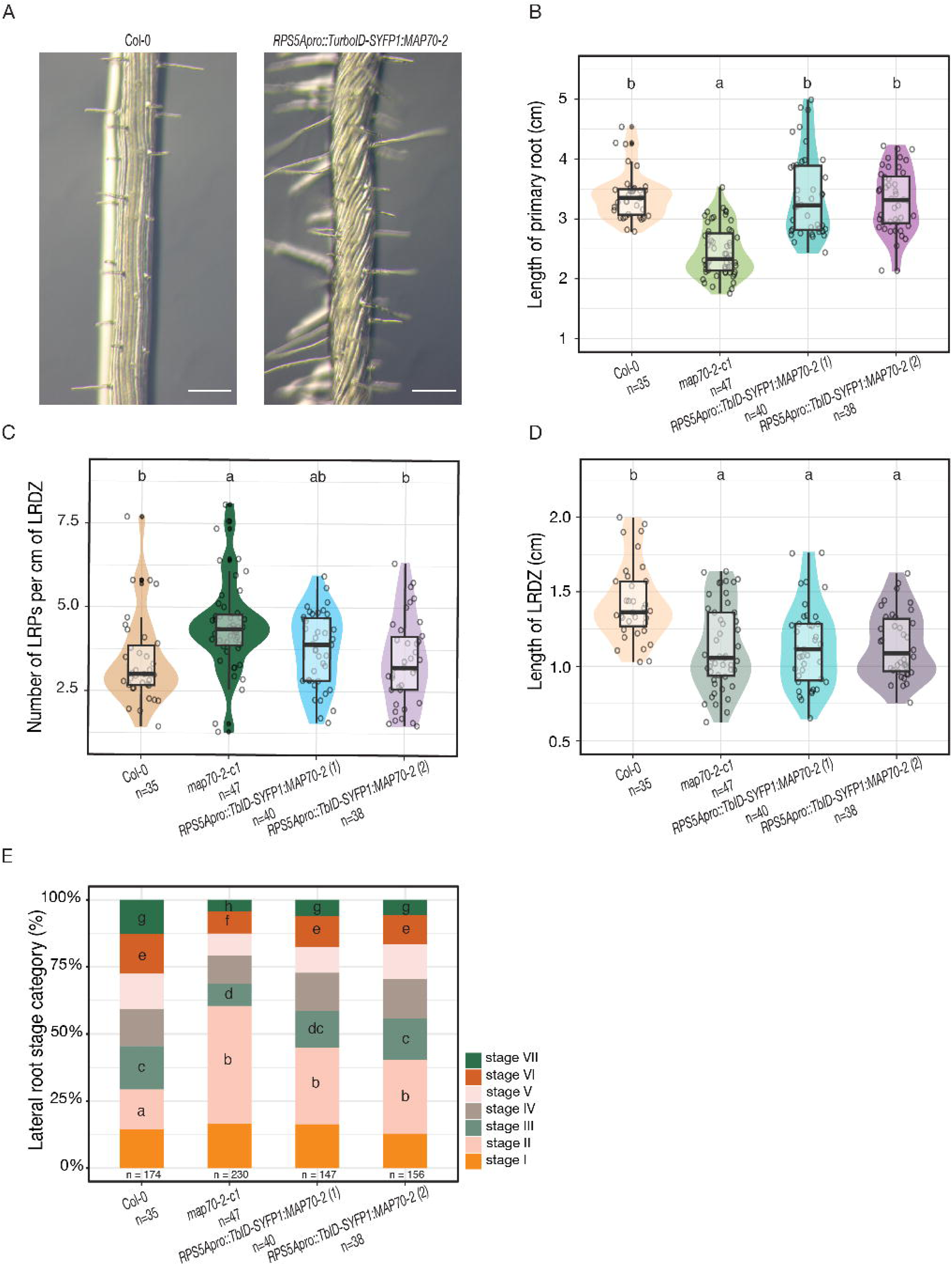
Overexpression of *MAP70-2* promotes left-handed root growth and right-handed epidermal cell file rotation. (A) Stereomicroscopic images showing root cell file rotations in 7-day-old Col-0 and the *RPS5Apro::TurboID-sYFP:MAP70-2* overexpression line, zoomed in to the elongation zone. Scale = 1 mm. (B) Comparison of primary root length between Col-0, *map70-2-c1* and two independent *RPS5Apro::TurboID-sYFP:MAP70-2* lines. Letters indicate significant differences (Kruskal-Wallis followed by Wilcoxon rank-sum tests, p *<* 0.05). (C) Comparison of LRP density between Col-0, *map70-2-c1* and two independent *RPS5Apro::TurboID-sYFP:MAP70-2* lines (pairwise Wilcoxon rank-sum tests with Benjamini-Hochberg correction,p < 0.05). (D) Comparison of LRP developmental stages between Col-0, *map70-2-c1* and two independent *RPS5Apro::TurboID-sYFP:MAP70-2* lines. Numbers indicate total primordia and seedlings analyzed. Different letters denote significant differences (Pearson’s Chi-squared test with Yates’ continuity correction, p< 0.05).

## Discussion

Lateral root development is complex process that occurs deep within the root, requiring precise coordination of molecular regulators to overcome the mechanical constraints imposed by the overlying cell layers. Previous studies have demonstrated that cytoskeletal reorganization is essential not only within the developing primordium, but also in the overlying endodermal cells that accommodate the growth of the new organ [33]. Although MAPs have been shown to be implicated in division plane positioning and primordium patterning [9,10,40,41], our findings identify MAP70-2 as a previously unrecognized regulator with organ-specific roles.

Our findings identify MAP70-2 as a previously uncharacterized regulator of cell division plane orientation during LR organogenesis, potentially via altering CMT organization. MAP70-2 binds MTs in vitro, its cellular localization depends on an intact MT cytoskeleton and MAP70-2 decorates the cortical division zones prior to cytokinesis, as well as spindles during divisions within the developing primordia (Figure 1E, Supplemental Figure 3A and B). Loss of *MAP70-2* resulted in increased numbers of stage II primordia, deformed LRP phenotype resulting from altered division plane orientation (Figure 2I), suggesting that MAP70-2 is required for proper establishment of division plane angles early during LR morphogenesis. Interestingly, MAP70-2 appears not to be required for division plane angle establishment in the primary root, as *map70-2-c1* roots had a similar organization compared to wild type, although the mutant roots are shorter due to a reduced meristematic zone (Figure 2C).

Our results support a role for MAP70-2 in regulating cell division plane organization during three-dimensional expansion growth within a tissue, possibly by affecting CMT structure, like what has been reported for MAP70-5 during xylem cell differentiation [32]. The phenotypes observed in *map70-2-c1* support a role for MAP70-2 as an integrator of mechanical constraints during 3D differential growth, required for proper establishment of division planes. A similar role was proposed for MAP70-5 in the endodermis, where it was reported to integrate mechanical signals, via CMT organization, to accommodate the expansion growth in the underlying XPP and primordium [33]. Our results also suggest that the LR phenotypes for *map70-2-c1* (Figure 2) and *map70-5-c1* [33] depend on the presence of MAP70-5 or MAP70-2 in the neighboring tissues, respectively, since the LR morphogenesis defects were complemented in the *map70-2/5* double mutant (Figure 4F and G). In contrast, the LR density was still enhanced and primary root length was still reduced in the double mutant (Figure 4 C and B).

The functional interplay between MAP70-2 and MAP70-5 highlights a broader division of labor within the MAP70 family. Whereas MAP70-5 operates primarily in differentiating cell types, including the endodermis and xylem [32,33], MAP70-2 acts in proliferative tissues where MT arrays are highly dynamic. Promoter-swapping experiments revealed a partial redundancy in primary root elongation, but non-overlapping functions during LR formation (Figure 3). This distinction may arise from differences in their biochemical properties or from the developmental contexts in which they operate, dividing primordium cells versus overlying, differentiated endodermal cells. Overexpression of MAP70-2 under the *RPS5A* promoter caused left-handed root twisting and right-handed epidermal cell-file rotation (Figure 5A), indicative of altered CMT alignment [34]. Such helical growth has been associated with excessive MT stabilization or bundling [34,42,43], suggesting that elevated MAP70-2 levels restrict the dynamic reorientation of cortical arrays. The concomitant accumulation of stage II primordia, despite a normal initiation frequency, supports the notion that a hyper-stabilized cytoskeleton impedes the structural remodeling required for LRP progression. Together with the *map70-2* loss-of-function phenotypes, these results suggest that MAP70-2 dosage balances MT stability and flexibility: reduced levels lead to disorganized arrays, while overexpression locks them in a rigid configuration.

Drawing parallels with MAP70-5 function in the overlying endodermis, we propose that MAP70-2 and MAP70-5 operate in complementary mechanosensory domains during LR formation. MAP70-2 acts within the primordium to organize MTs during formative divisions, whereas MAP70-5 functions in adjacent tissues to relieve mechanical resistance to primordium growth, possibly via altering the cell wall composition. This coordinated activity would ensure that differential growth within the root is matched by appropriate cytoskeletal adaptation in neighboring cells. In this model, MAP70 proteins act as integrators of local mechanical constraints and MT organization, channeling encountered mechanical constraints to support LR morphogenesis.

## Materials and methods

### Plant materials and manipulation

*Arabidopsis thaliana* Columbia ecotype (Col-0) was used. Seeds were surface sterilized using chlorine gas and placed on ½ Murashige and Skoog (MS) pH 5.8 medium containing 0,8 % Phyto Agar (Duchefa Biochemie). Following stratification (4 °C in the dark for at least 48 hours), seedlings were grown vertically at 22 °C under constant light or long-day conditions (120 µmol m⁻² s⁻¹). *Agrobacterium tumefaciens-*based plant transformation was carried out using the floral dip method [44]. All plant lines examined were homozygous, unless otherwise indicated. Homozygosity was determined by the presence of the Fast Red cassette, antibiotic resistance, verification of fluorescent fusion proteins under the microscope, and/or PCR. Besides the below-mentioned created vectors and plant lines, this study uses the LRP membrane marker line *pGr179-LBD16pro::3xmCherry:SYP122* [33], the LRP actin marker *LBD16pro::ABD2:GFP*, the microtubule cytoskeleton marker line *pGr179-Col-0-XPPpro::mVENUS:MBD* [38] and the membrane marker line *UBQ10pro::EYFP: NPSN12* (*WAVE131Y*) [35].

### Construction of vectors and generation of transgenic lines

For *pEN-2-gMAP70-2-3*, the genomic sequence of *MAP70-2* (At1g24764) was amplified and recombined into pENTR2-3 using BP CLONASE II (www.thermofisher.com).

To generate *pFR-MAP70-2pro::CITRINE:gMAP70-2 and pFR-MAP70-2pro::TurboID-sYFP:gMAP70-2*, we cloned *TurboID-SYFP1* into *pDONR221* using infusion HD cloning [45]. Finally, all fragments, *pEN-4-MAP70-2-1*, *pEN1-CITRINE-2* or *pEN1-TurboID-sYFP-2* and *pEN-2-gMAP70-2-3* were recombined into *pFastRed-3xGW* using LR CLONASE II plus (www.thermofisher.com).

The following plasmids were generated using GreenGate modular cloning [46]. For *pFR-LBD16pro::TurboID-sYFP:MBD, pFR-MAP70-2pro::GFP-NLS:GUS, pFR-RPS5Apro::TurboID-sYFP:MAP70-2, pFR-RPS5Apro::TurboID-sYFP:GFP, pFR-MAP70-2pro::TurboID-sYFP:MAP70-5* the following modules were assembled in *pFASTR-AG*: (A) *MAP70-2pro* or (A) *LBD16pro* or (A) *RPS5Apro*, (B) *TurboID-sYFP* or (B) *SV40 NLS*, (C) *gMAP70-5* or (C) *GUS*, (D) *D-dummy*, (E) *At1g04880* terminator, (F*) F-dummy*. The *MAP70-2pro* module was obtained by PCR of a 2802 bp fragment upstream of the ATG of *MAP70-2*. The same method was used for *TurboID-sYFP* (1740 bp), and g*MAP70-5* (2398 bp). All the other modules are described in Lampropoulos et al., 2013. For creating the GUS marker lines, the following modules were combined in pFASTR-AG: (A) *MAP70-2pro*, (B) *SV40NLS*, (C) *GUS*, (D) *linker-GFP*, (E) *At1g04880* terminator, (F) *F-dummy*. All plasmids were sequenced before usage. Arabidopsis plants were transformed using the floral dip method [44]. For CRISPR/Cas9-mediated generation of *map70-2* mutants, we used *pFR-UBQ-CAS9-1xGW*. Three different sgRNA constructs were cloned, targeting different areas of MAP70-2, using a combination of GreenGate [46] and Gateway cloning. Primers for amplifying fragments used for cloning and sgRNAs used for generating *map70-2-c1* and *-c2* mutants are shown in Table 1. The genomic sequences of *map70-2-c1* and *map70-2-c2* were amplified and recombined into pDONR1-2 for sequence verification of the introduced deletions.

**Table 1.**
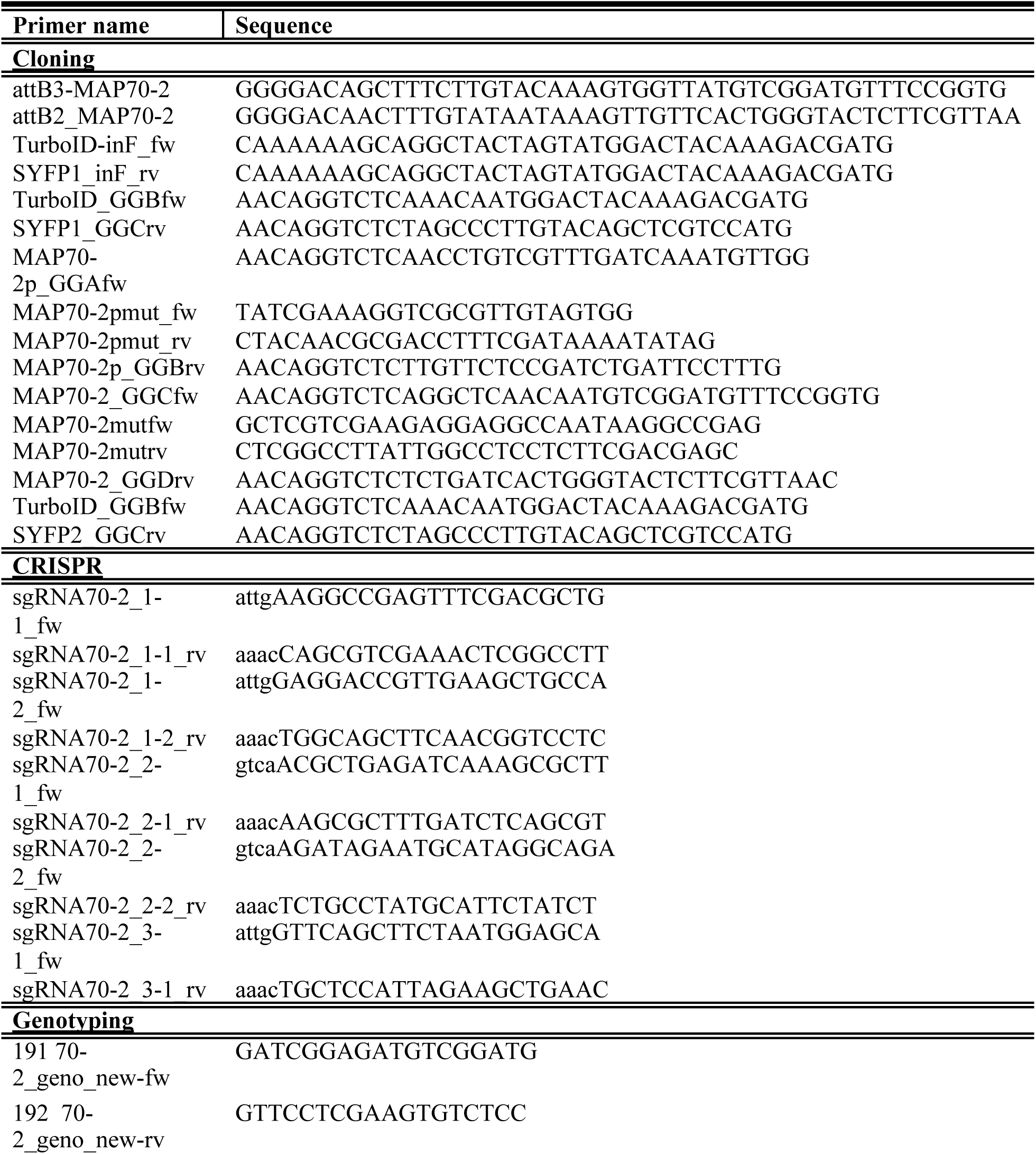
Primers used for cloning, CRISPR and genotyping.

### Oryzalin Treatment

To disrupt microtubule organization, seedlings were treated with the microtubule-depolymerizing agent oryzalin. Arabidopsis seedlings were grown vertically on half-strength Murashige and Skoog (½ MS) medium solidified with 0.8% agar under continuous light at 22°C. For chemical treatment, 5-day-old seedlings were transferred to liquid ½ MS medium containing 500 nM oryzalin (Sigma-Aldrich) or an equivalent volume of DMSO as a mock control. Seedlings were incubated in the treatment solution for 16 hours prior to imaging with gentle agitation to ensure uniform exposure. Oryzalin stocks were prepared in DMSO and stored at −20°C.

### GUS staining

Histochemical GUS staining was performed to assess reporter gene expression in 7-day-old seedlings. To fix and remove chlorophyll, seedlings were harvested and incubated in cold 90% acetone and incubated for 20 min. Later, seedlings were transferred into GUS staining buffer consisting of 100 mM sodium phosphate buffer (pH 7.2), 0.2% Triton X-100, 0.75 mM potassium ferrocyanide, 0.75 mM potassium ferricyanide, Dimethylformamide (DMF), and 2 mM 5-bromo-4-chloro-3-indolyl-β-D-glucuronide (X-Gluc, Thermo Scientific) as the substrate. Samples were vacuum infiltrated for 20 minutes and then incubated at 37°C up to 16 hours in the dark. Following staining, tissues were cleared in a graded ethanol series ( 95%, 70%, 50%, 30% 10%) for 5 min each to enhance visualization. Stained samples were stored in 50% glycerol and stored in 4°C until processing.

### Protein purification

To generate 6×His–mGFP–MAP70-2, the *MAP70-2* fragment was PCR-amplified from Arabidopsis cDNA and inserted into the pColdTEV–6×His–mGFP–MAP70-5 vector [32] in place of the *MAP70-5* sequence using In-Fusion HD cloning kit (Takara). The 6×His-mGFP-MAP70-2 construct was transformed into the *Escherichia coli* BL21-CodonPlus (DE3)-RILP competent cells (Agilent Technologies). The competent cells were incubated at 37 °C until the OD_600_ reached 0.5 and then further incubated at 15°C overnight with 0.1 mM isopropyl-β-D-thiogalactopyranoside. Pellets of *E. coli* were collected, resuspended in His elution buffer [50 mM Tris-HCl (pH 7.5), 500 mM NaCl, 20 mM imidazole], and sonicated. The cell lysate was centrifuged at 3000 × *g* for 2 min, and the supernatant was incubated with Ni Sepharose 6 Fast Flow resin (GE Healthcare) at 4 °C for 1 h. Recombinant proteins were eluted with 500 µl of His-elution buffer [50 mM Tris-HCl (pH 7.5), 500 mM NaCl, and 500 mM imidazole]. Imidazole was removed from the eluate using a PD MiniTrap G-25 column (GE Healthcare) with PIPES buffer [10 mM PIPES (pH 6.9), 50 mM DTT, 5% sucrose, and 500 mM NaCl].

### Preparation of rhodamine-labeled microtubules

Tubulin was purified from porcine brain using the high-molarity PIPES procedure [47] and labeled with X-rhodamine as described previously[32]. Rhodamine-labeled and unlabeled tubulins were mixed at a molar ratio of 1:20. Microtubules were polymerized in BRB80 buffer (80 mM K-PIPES, pH 6.8, 1 mM MgCl₂, 1 mM EGTA) containing 1 mM GTP, 4 mM MgCl₂, and 5% DMSO at 37 °C for 30 min. The polymerized microtubules were stabilized in BRB80 supplemented with 10 µM taxol, pelleted by ultracentrifugation (230,000 × *g*, 25 °C), and resuspended in BRB80 containing 10 µM taxol.

### Microtubule binding assay

Flow chambers were assembled by attaching a glass coverslip on a glass slide with two double-stick tape aligned in parallel with ∼3 mm separation. A glass slide flow chamber was incubated sequentially with the following reagents: 0.5mg/ml PLL-g-PEG-biotin for 5 min, 0.1 mg/ml streptavidin for 5 min, blocking buffer (1.0 mg/ml casein, 10 mg/ml BSA, and 10 µM Taxol) for 2 min, and MAP70-2-microtubule mixture (X-rhodamine-labeled biotinylated microtubules and 5 µM mGFP-MAP70-2) for 10 min. The chamber was flushed with assay buffer before imaging. Microtubule binding of mGFP-MAP70-2 was imaged using a Nikon ECLIPSE Ti2-E TIRF microscope with an Apo TIRF 100× oil objective lens (1.49 NA, Nikon), an EMCCD camera (iXon3 888, Andor Technology), and an LUA-S4 laser unit (Nikon).

### Microscopy

Live cell imaging *was* performed with a Leica TCS SP8X-MP, TCS SP8-MP-DIVE or Leica Stellaris WLL-plus confocal microscope equipped with a resonant scanner (8 kHz), a 63x, NA = 1.2 water immersion objective. For the excitation of GFP, YFP, or CITRINE a Chameleon Vision II tuned to 960 nm was used for multiphoton excitation. For detection, non-descanned super-sensitive photon-counting hybrid detectors (HyD), operated in the photon-counting mode, were used. For imaging of *MAP70-2pro::CITRINE:MAP70-2* and *pGr179-LBD16pro::3xmCherry:SYP122*, we used 960nm 2P-excitation and 500-550 nm detection for CITRINE and 590-650 nm for mCherry using a 4-TUNE non-descanned detector. The acquired z-stacks in this study were taken in step sizes ranging from 0.3-0.5 µm. For long-term imaging, time-lapse data were acquired every 15-20 min for up to 13 h. Epidermal cell file rotation and whole seedling GUS staining images were taken with a Leica Flexcam C5 Microscope Camera.

### Phenotyping *map70-2* mutants

Roots were scanned from the back of the plates using an Epson Perfection V600 Photo Scanner, and primary root length was measured from the root tip to the base of the hypocotyl in seven-day-old Col-0 and *map70-2* mutants. To quantify LRP density in the three genotypes along the LRDZ, the seven-day-old seedlings were cleared as published Voß et al., 2015. Experiments were repeated three times. Col-0 and *map70-2* mutants were grown on different plates, made from the same batch of plant agar. Upon clearing, seedlings were mounted in 50% glycerol and staged using Leica DM4 upright microscope equipped with differential interference contrast. Staging was always initiated from the first stage I LRP and proceeded until the first emerged LRP. Using a permanent marker, we would indicate on the cover slip of the sample the first stage I LRP and the first emerged LRP to define the length of the LRDZ. Marked slides would then be scanned and the LRDZ length would be measured using the SmartRoot plugin in FIJI. To visualize the cell outlines, *map70-2-c1* plants were crossed with the *UBQ10pro::EYFP:NPSN12* cell membrane marker line, and homozygous F_3_ plants were imaged and staged from stage I until stage III (Leica TCS SP8-MP-DIVE 960 nm excitation, 500 – 600nm detection).

### Meristem-size analysis

The length of the meristematic zone was measured based on the number of cortex cells, following the definition established by Dello Ioio et al.[49]. Specifically, the meristematic region was considered to extend from the quiescent center (QC) to the first cortex cell that was at least twice as long as the one directly before it, marking the transition from isodiametric to elongated cells.

### Analysis of division plane orientation

To analyze the orientation of cell divisions during lateral root development, 5-day-old *Arabidopsis thaliana pFR-LBD16pro::TurboID-sYFP:MBD* seedlings were grown vertically on ½ MS and 0.8% agar plates with 1% sucrose for under the same conditions as mentioned above. At 4 days after germination, seedlings were transferred to imaging chambers containing ½ MS and 1% plant agar with 1% sucrose. Live imaging was performed using a confocal laser scanning microscope Leica TCS SP8-MP-DIVE up to 13 h. Time-lapse imaging was carried out overnight at 15-20 min intervals using a 63× water-immersion objective. The angle was measured between the transverse cell walls and the adjacent longitudinal cell wall of the developing LRP. Division planes were determined by tracking mitotic events and classified as anticlinal, periclinal, based on their orientation relative to the radial axis of the root. Division-plane orientation angles were measured using the Angle tool in Fiji.

### Data visualization and statistics

All experiments were performed at least three times. Sample size (n) for each plant line and treatment is denoted in the figures. Statistical comparisons (two-way ANOVA, Wilcoxon rank-sum test, Pearson’s Chi-squared test) and graphing were made with RStudio Version 2023.12.1+402. Adjustments and figure assembly were performed with Adobe Illustrator 28.7.1.

### Image processing

The acquired microscopy images were processed using FIJI (v2.1.0/1.53k, https://fiji.sc/).

## Acknowledgements

Work in the Vermeer lab was funded by the Swiss National Science Foundation (project nos.: 157524, 197568 and 231162 awarded to JEMV) and by the University of Neuchâtel. Work in the Oda lab was supported by MEXT KAKENHI (Grant no. 25H02364 to YO) and JSPS KAKENHI (Grant nos. 24K02042, 23K18126, 24H00056 to YO, and 23K05801 to TS)

---

**Supplementary Figure 1. Different deletions introduced in *map70-2-c1* and *-c2*.** Schematic representation of the gene model of *MAP70-2* and the position of the different sgRNAs used to generate *map70-2* mutants. The bottom panel displays the generated deletions, all of which lead to premature stop codons, indicated by the underlined pink letters.

**Supplementary Figure 2.** Schematic representation of the quantification of cell division orientations in growing LRP. Cell divisions were analyzed during cytokinesis, specifically at the stage of cell plate formation. The orientation of each division was measured relative to the growth axis of the primordium.

**Supplementary Figure 3. Complementation of the *map70-2* mutant with genomic *MAP70-2* restores wild-type root phenotypes in 7-day-old *Arabidopsis* seedlings.**

(A) Representative seedlings of 7-day-old seedlings expressing *MAP70-2pro::CITRINE:MAP70-2* in Col-0 and in the *map70-2c-1* mutant background. Scale bar = 1 cm.

(B) Quantification of primary root length and LRDZ (D) using Student‘s t-test. There was no significant difference between *MAP70-2pro::CITRINE:MAP70-2/* Col-0 and the complemented *map70-2-c1* (p<0.05).

(C) The Wilcoxon rank-sum test revealed no significant difference in LRP density between Col-0 background and the complemented *map70-2-c1* (p<0.05).

(D) Quantification LRDZ using Student‘s t-test. There was no significant difference between *MAP70-2pro::CITRINE:MAP70-2/* Col-0 and the complemented *map70-2-c1* (p<0.05).

(E) Bar plot showing the distribution of developmental stages of LRPs. No significant differences (n.s) were detected between *MAP70-2pro::CITRINE:MAP70-2* lines in Col-0 and *map70-2-c1* across all measured parameters, indicating successful complementation of the *map70-2* mutant phenotype. Sample sizes (n) are indicated below each graph.

**Supplementary Figure 4. CITRINE:MAP70-2 localization is dependent on intact microtubules and MAP70-2 does not regulate cell division plane orientation in the root apical meristem.** (A and B) Confocal images of root tips and stage I or stage II LRP expressing MAP70-2*pro::CITRINE:MAP70-2*. (A) CITRINE:MAP70-2 localization after 16 h of mock (DMSO) or (B) after oryzalin (500 nM) treatment. Scale bar = 20 µm. (C and D) Confocal images of the primary root meristem visualized using *UBQ10pro::EYFP:NPSN12* (W131Y) in control (C) and *map70-2-c1* mutant (D) seedlings, showing comparable meristem organization. Scale bar = 20 µm.

