## Supplementary figures and images for "A MICROTUBULE ASSOCIATED PROTEIN is required for division plane orientation during 3D-differential growth within a tissue"

### Supplemental Figure S1

#### *MAP70-2 (AT1G24764.3)*

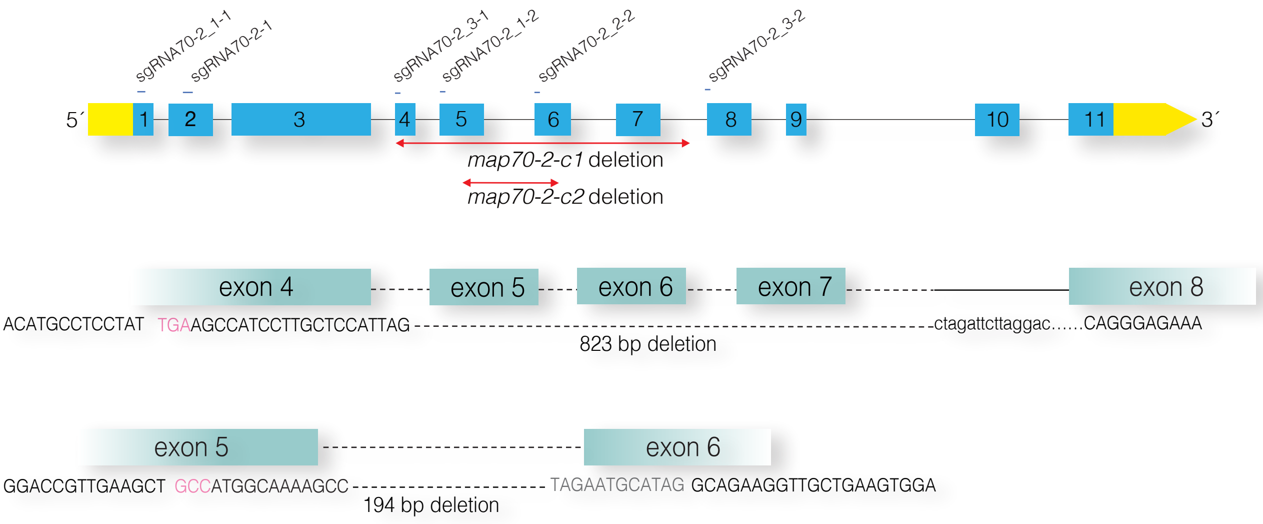

### Supplemental Figure S2

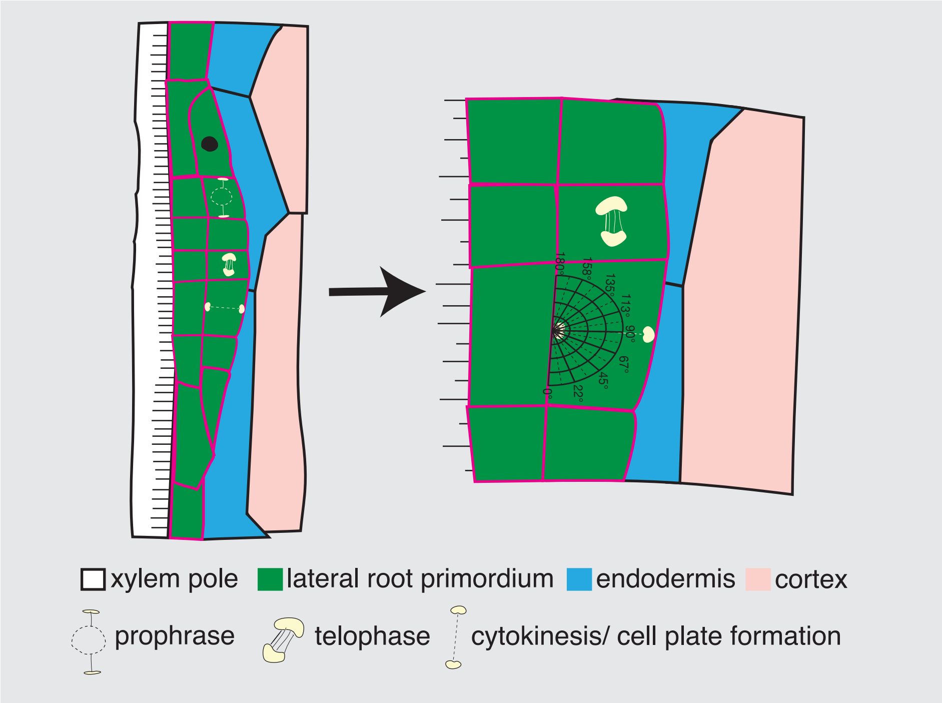

### Supplemental Figure S3

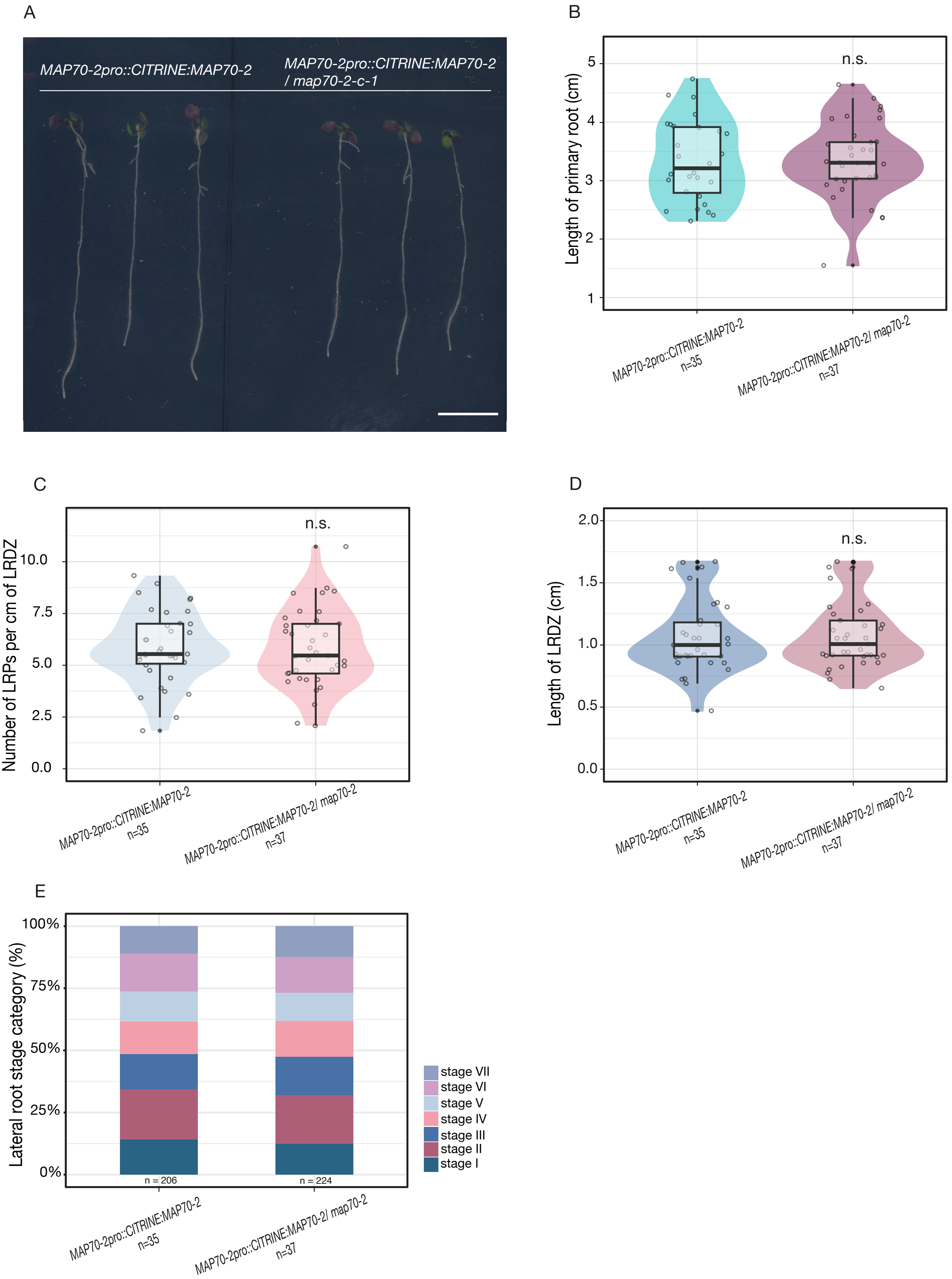

### Supplemental Figure S4

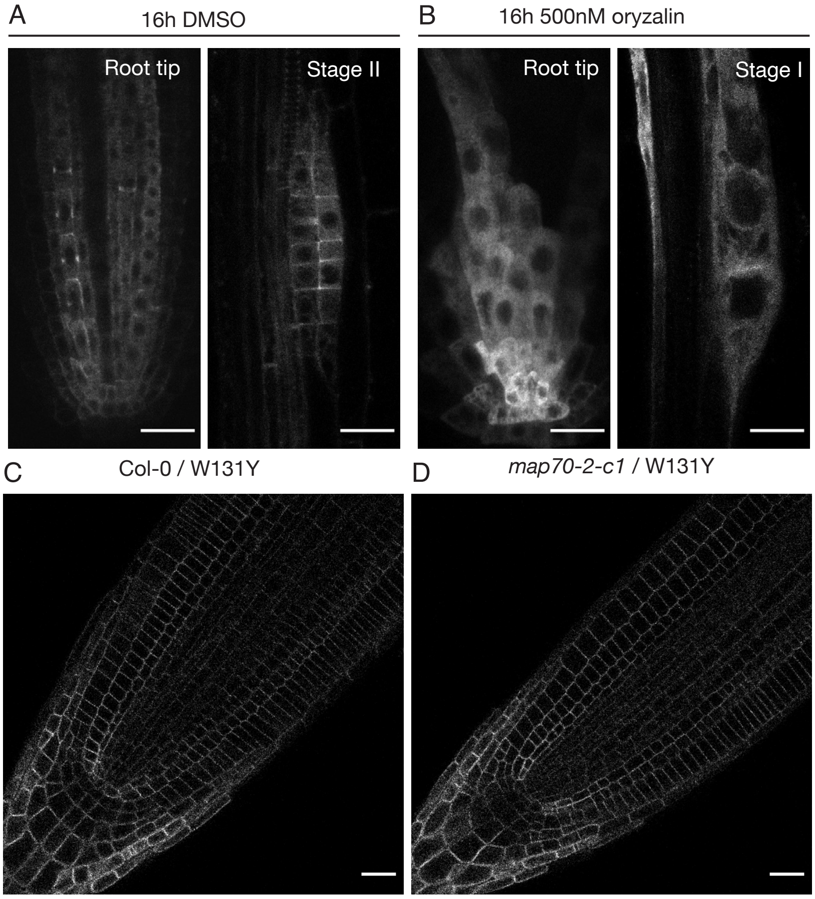
